# Abrogation of RAB27A expression transiently affects melanoma cell proliferation

**DOI:** 10.1101/2019.12.27.889329

**Authors:** Dajiang Guo, Kimberley A. Beaumont, Danae M. Sharp, Goldie Y. L. Lui, Wolfgang Weninger, Nikolas K. Haass, Shweta Tikoo

**Author notes:** Corresponding authors: Nikolas K. Haass, The University of Queensland, Diamantina Institute, Translational Research Institute, 37 Kent St, Woolloongabba, QLD 4102, Australia., Shweta Tikoo, The Centenary Institute of Cancer Medicine and Cell Biology, Locked Bag 6, Newtown, NSW, 2042, Australia. These authors contributed equally to this work (Nikolas K. Haass, Shweta Tikoo).

## Abstract

The role of the small GTPase RAB27A as an essential melanosome trafficking regulator in melanocytes is well-accepted. A decade ago, RAB27A was identified as a tumor dependency gene that promotes melanoma cell proliferation. RAB27A has since been linked to another propeller of cancer progression: exosome secretion. We have recently demonstrated that RAB27A is overexpressed in a subset of melanomas. High RAB27A gene and protein expression correlates with poor prognosis in melanoma patients. Mechanistic investigations revealed that the generation of pro-invasive exosomes was RAB27A-dependent and, therefore, silencing RAB27A reduced melanoma cell invasion *in vitro* and *in vivo*. However, previous studies have implicated RAB27A to be involved in both proliferation and invasion of melanoma cells. In this study, we demonstrate that the effects of abrogating RAB27A expression on proliferation are temporary, in contrast to the previously reported persistent effects on tumor invasion and metastasis. Therefore, we assist in the dissection of the short-term versus long-term effects of RAB27A knockdown on melanoma cell proliferation, invasion, and metastasis. We believe that our findings provide novel insights into the effects of RAB27A blockade.

**Significance:** RAB27A is known to serve as an essential regulator for melanosome trafficking. However, to date its role in melanoma biology has not been completely deciphered. While there are consistent independent reports on the pro-invasive effects of RAB27A, there are conflicting data on its impact on cell proliferation. Here we show that indeed abrogation of RAB27A does reduce cellular proliferation; however, this effect is only transient, while the impact on invasion as reported previously is persistent. This finding offers an explanation for the apparent contradiction in the literature and provides a deeper understanding of RAB27A function in melanoma cell biology.

## Main Text

The small GTPase RAB27A is a well-recognized regulator of intracellular signaling and mediator of melanosome trafficking in melanocytes (Strom, Hume, Tarafder, Barkagianni, & Seabra, 2002). Past studies have linked RAB27A to promoting melanoma growth and metastasis. Utilizing a computational algorithm termed CONEXIC (copy number and expression in cancer), RAB27A was identified as a tumor dependency gene in melanoma (Akavia et al., 2010). The authors went on to validate the role of RAB27A in melanoma cell proliferation by demonstrating that shRNA-mediated knockdown of RAB27A in short-term cultures with elevated RAB27A expression levels led to reduced cell growth as against the control cultures with low RAB27A expression levels which showed no reduction in cellular proliferation (Akavia et al., 2010). Furthermore, Peinado and colleagues have demonstrated that abrogation of RAB27A expression leads to reduced exosome secretion in melanoma cells and thereby suppresses melanoma metastasis. They further established that the pro-tumoral function of RAB27A is mediated by tumor cell-derived exosomes that home to the bone marrow and educate the bone marrow progenitor cells towards a more vasculogenic phenotype, thus ultimately assisting in the creation of a pre-metastatic niche (Peinado et al., 2012). We have recently dissected the role of RAB27A in melanoma invasion and metastasis (Guo et al., 2020; Guo, Lui, et al., 2019; Guo, Tikoo, & Haass, 2019). Our study revealed that both *RAB27A* RNA and protein expression levels correlated with poor survival in melanoma patients (Guo, Lui, et al., 2019). Moreover, the abrogation of RAB27A expression in melanoma cells led to a marked reduction in tumor invasion and metastasis both *in vitro* and *in vivo* (Guo, Lui, et al., 2019). We found that this was caused by the reduction in the secretion of pro-invasive exosomes, even though the total number of secreted exosomes remained the same (Guo, Lui, et al., 2019). These findings identified RAB27A as a key cancer regulator, as well as a potential prognostic marker and plausible therapeutic target for the treatment of advanced melanoma.

Various studies to date have implicated RAB27A in promoting cell proliferation and tumor growth in many types of cancers, including melanoma (Akavia et al., 2010; Peinado et al., 2012), bladder (Liu et al., 2017), breast (Bobrie et al., 2012) and pancreatic cancer (Li, Jin, Huang, Tang, & Huang, 2017). A recent study demonstrated that abrogation of *RAB27A* expression caused a reduction in RAB27A-mediated exosome secretion, which resulted in a cell proliferation defect (Li et al., 2019). In concordance with these studies, we have also found a proliferation defect post abrogation of *RAB27A* expression in melanoma cells. However, we show that this defect in proliferation is transient (this study), while the defects in cell invasion and metastasis were persistent (Guo, Lui, et al., 2019). This suggests that the effects of *RAB27A*-knockdown on melanoma cell proliferation may be more complex than initially anticipated.

To understand the role of RAB27A in melanoma cell proliferation, we chose two endogenously RAB27A-high melanoma cell lines, namely WM164 and WM983C (Figure 1A, B) for abrogation of *RAB27A* expression. *RAB27A*-knockdown (KD) employing transduction with two different shRNA-expressing lentiviral particles initially caused modest cell death and more rounded cell morphology (Figure 1C, Supplementary Figure 1A). This was not a transduction artifact as cells transduced with the non-targeting shRNA showed no such effect. However, after three passages (approximately 4 weeks), the surviving WM164 and WM983C *RAB27A*-KD cells started to regain normal morphology (Figure 1D, Supplementary Figure 1B and F-actin structure (Figure 1E). Immunoblotting showed that the surviving *RAB27A*-KD cells had maintained RAB27A abrogation (Figure 1F, G).

**Figure 1.**
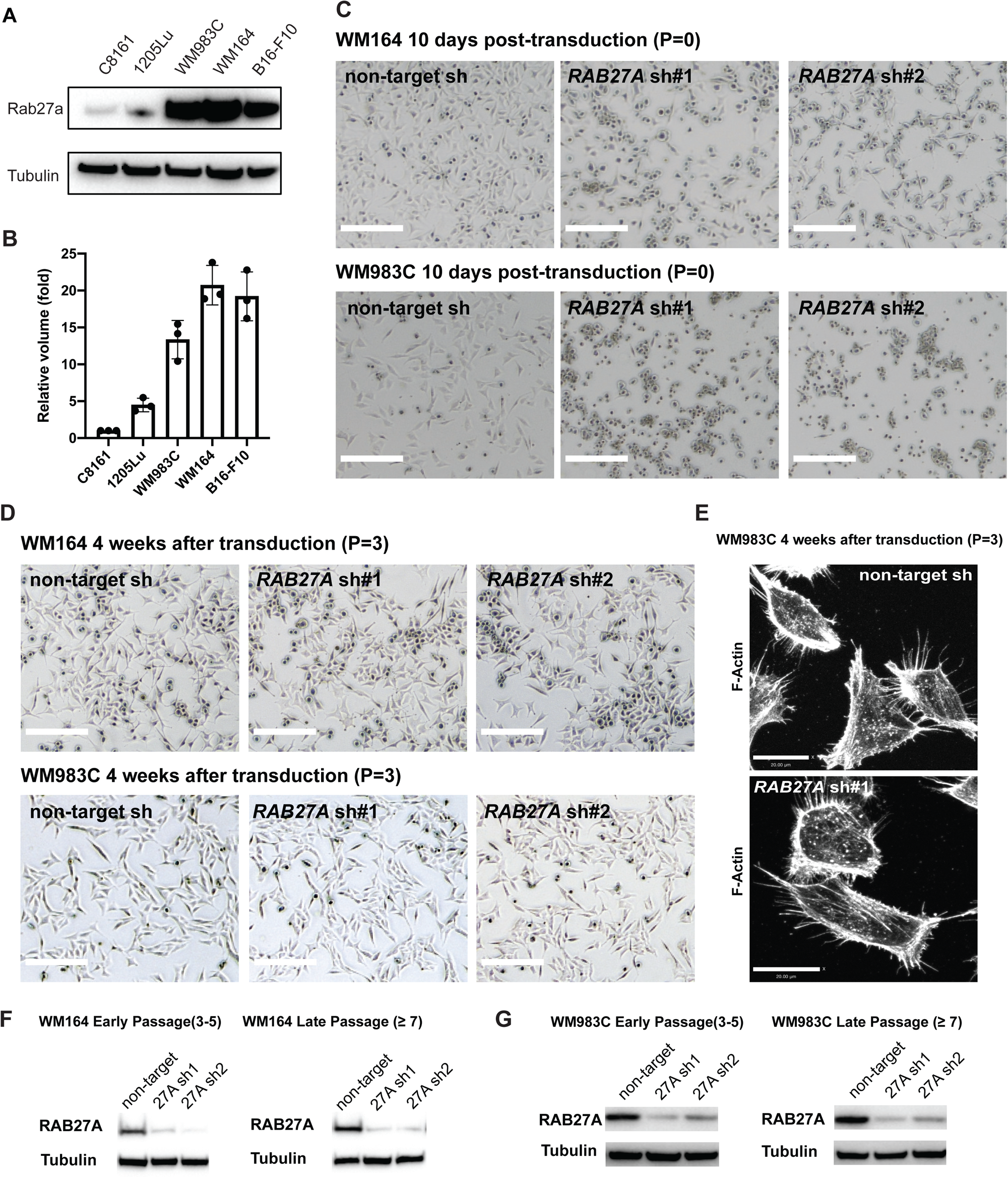
*RAB27A*-knockdown in RAB27A-high melanoma cell lines causes cell death and morphology alteration only in early passages. **(A)** Western blot showing RAB27a protein levels in five melanoma cell lines as indicated. Representative of 3 independent experiments. **(B)** Quantification of RAB27a protein levels in five melanoma cell lines as indicated, based on the protein band volume from Western blotting. Protein volumes are normalized to C8161. Error bars represent the mean ± SEM of 3 independent experiments. **(C)** Phase contrast images of WM164 and WM983C melanoma cells approximately 10 days after transduction with *RAB27A* shRNA or control lentivirus and 8 days after selection with puromycin. *RAB27A* shRNA KD cells are at Passage 0 after transduction. Note that all untransduced cells were dead after 2-3 days of puromycin selection. Scale bars = 100μm. Representative of 3 independent experiments. **(D)** Phase contrast images of WM164 and WM983C melanoma cells approximately 4 weeks (3 passages) after transduction with *RAB27A* shRNA or control lentivirus Scale bars = 100μm. Representative of 3 independent experiments. **(E)** Confocal extended focus image of F-actin staining in WM983C cells transduced with *RAB27A* shRNA or non-targeting shRNA (passage 3 after transduction). Representative of 3 independent experiments. **(F, G)** Western blots showing the level of RAB27A protein in WM164 (**F**) and WM983C cells (**G**), non-targeting shRNA control (non-target), *RAB27A* shRNA1 (27A sh1) and *RAB27A* shRNA#2 (27A sh2) (early = passage 3-5 after transduction, late ≥ passage 7). Representative of 3 independent experiments.

Early passage (passages 3-5 after transduction) WM164 and WM983C *RAB27A*-KD cells showed proliferation and colony-forming defects (Figure 2A, B, C) that were gradually lost at high passage numbers after transduction (≥7 passages; Figure 2D, E, F). This was not due to reverting of RAB27A expression, as RAB27A protein levels remained low in the high passage shRNA expressing cells (Figure 1E, F).

**Figure 2.**
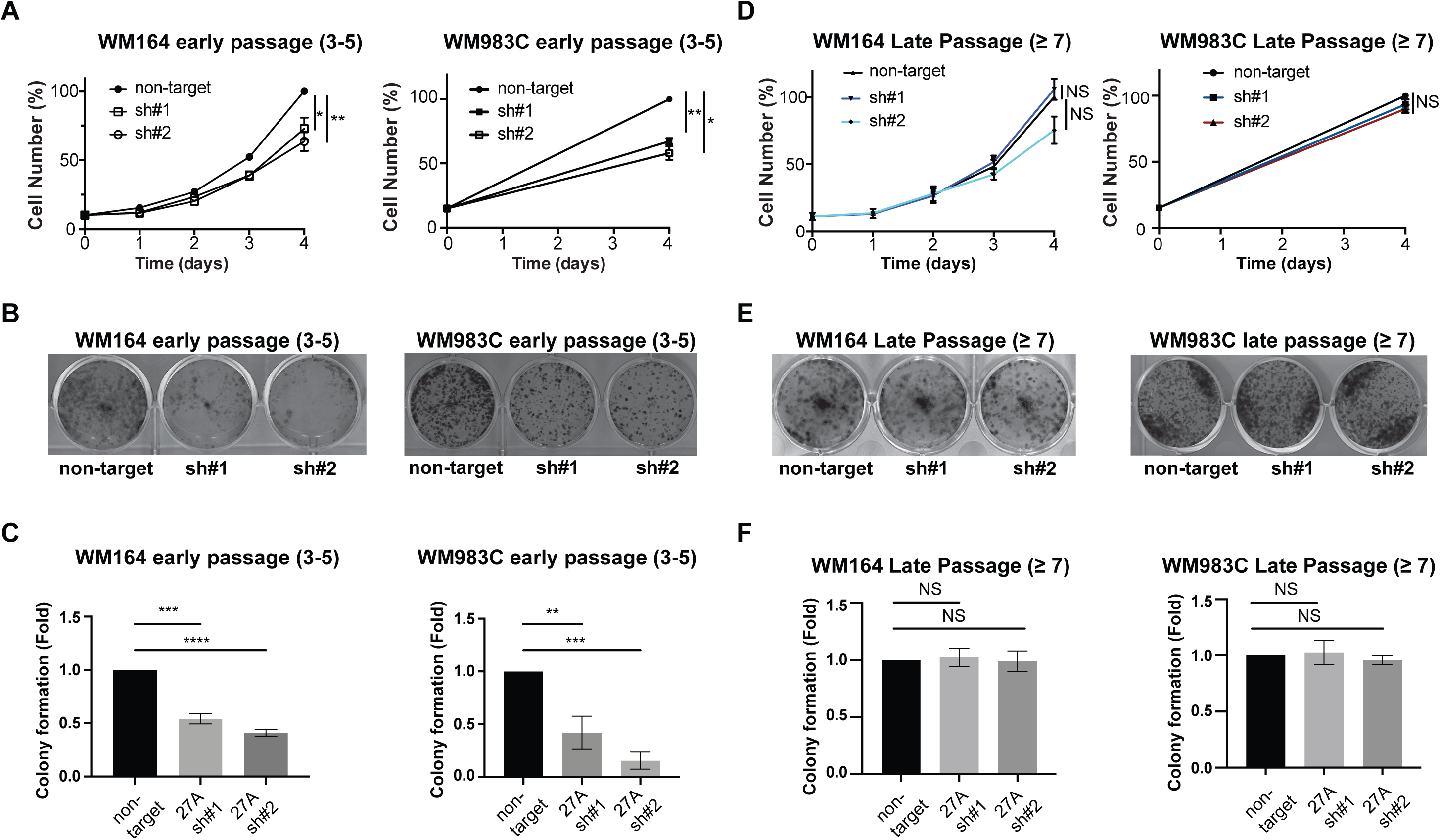
*RAB27A*-knockdown in RAB27A-high melanoma cell lines reduces proliferation and colony formation potential only in early passages. **(A)** Normalized cell counts over the indicated days in early passage (p3-5) WM164 or WM983C cells expressing the non-targeting shRNA control, *RAB27A* shRNA#1 or *RAB27A* shRNA #2. Error bars represent the mean ± SEM of 3 independent experiments. Non-targeting control was compared to the other samples via ANOVA and a Dunnett’s test at day 4. **(B)** Colony-forming assay of early passage (p3-5) WM164 or WM983C cells expressing the non-targeting shRNA control, *RAB27A* shRNA#1 or *RAB27A* shRNA #2. Representative of 3 independent experiments. **(C)** Quantification of colony formation of early passage (p3-5) WM164 or WM983C cells expressing the non-targeting shRNA control, *RAB27A* shRNA#1 or *RAB27A* shRNA #2. Error bars represent the mean ± SEM of 3 independent experiments. Non-targeting control was compared to the other samples via ANOVA and a Dunnett’s test. **(D)** Normalized cell counts over the indicated days in late passage (passage ≥ 7) WM164 cells expressing the non-targeting shRNA control, *RAB27A* shRNA#1 or *RAB27A* shRNA #2. Error bars represent the mean ± SEM of 3 independent experiments. Non-targeting control was compared to the other samples via ANOVA and a Dunnett’s test and no significance was detected. **(E)** Colony-forming assay of late passage (≥ 7 passage) WM164 and WM983C cells expressing the non-targeting shRNA control, *RAB27A* shRNA#1 or *RAB27A* shRNA #2. Representative of 3 independent experiments. (**F**) Quantification of colony formation of late passage (≥ 7 passage) WM164 or WM983C cells expressing the non-targeting shRNA control, *RAB27A* shRNA#1 or *RAB27A* shRNA #2. Error bars represent the mean ± SEM of 3 independent experiments. Non-targeting control was compared to the other samples via ANOVA and a Dunnett’s test. NS, not significant.

Because the normalization of proliferation and colony-forming defects could not be attributed to the recovery of *RAB27A* expression, we anticipated compensation by other members of the RAB GTPase family to be responsible for these observations. As it has been reported that RAB27B can compensate for the loss of RAB27A (Barral et al., 2002), we analyzed the expression of *RAB27B* mRNA via quantitative real-time PCR. We found that the expression of *RAB27B* in late passage *RAB27A*-KD cells was not significantly elevated compared to the control cells (Supplementary Figure 1C). Thus, compensation for the loss of RAB27A in late passage cells could not be conclusively attributed to an increase in *RAB27B* expression.

To determine the effect of the loss of RAB27A on survival and proliferation in a melanoma cell line endogenously expressing low levels of RAB27A protein, we performed *RAB27A*-KD in RAB27A-low C8161 cells (Figure 1A, B). Western blotting confirmed that *RAB27A* shRNA-expressing cells had substantially reduced RAB27A (Figure 3A). *RAB27A*-KD did not affect cell morphology, cell death, or proliferation in C8161 cells (Figure 3B, C). This indicates that RAB27A-low melanoma cells do not rely on RAB27A for proliferation or survival. *RAB27A*-KD (Figure 3A) in a melanoma cell line that has intermediate levels of RAB27A (1205Lu, Figure 1A, B) did not cause cell death (Figure 3B, Supplementary Figure 1A, B). However, similar to WM983C and WM164, *RAB27A*-KD in 1205Lu cells led to a reduced proliferation phenotype (Figure 3C).

**Figure 3.**
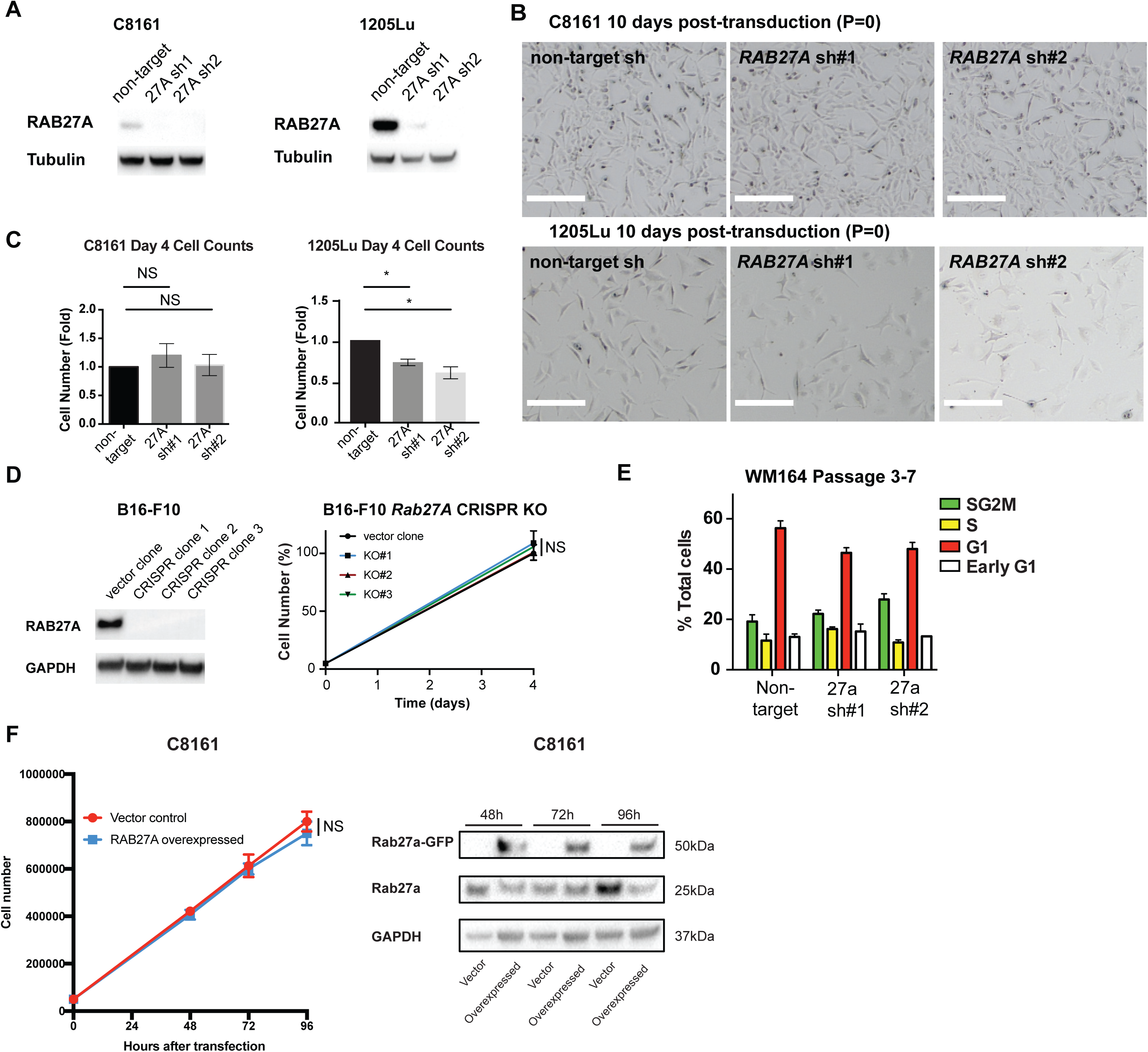
*RAB27A*-knockdown does not alter *RAB27B* expression, cell cycle profile as well as proliferation and colony formation in RAB27A-low melanoma cell lines. **(A)** Western blots showing the level of RAB27A protein in early passage (passage 3-5 after transduction) C8161 and 1205Lu, non-targeting shRNA control, *RAB27A* shRNA#1 (27A sh1) and *RAB27A* shRNA#2 (27A sh2) cells. Representative of 3 independent experiments. **(B)** Phase contrast images of C8161 and 1205Lu melanoma cells approximately 10 days after transduction with *RAB27A* shRNA or control lentivirus and 8 days after selection with puromycin. Scale bars = 100μm. Representative of 3 independent experiments. **(C)** Normalized cell counts over the indicated days in early passage (passage 3-5 after transduction) C8161 and 1205Lu cell lines expressing the non-targeting shRNA control, *RAB27A* shRNA#1 or *RAB27A* shRNA #2. Error bars represent the mean ± SEM of 3 independent experiments. Non-targeting control was compared to the other samples via ANOVA and a Dunnett’s test. No significance was detected in C8161 cell line comparisons. (**D)** Western blots showing the levels of Rab27A (left) and proliferation assay (right) of B16-F10 empty vector control, *Rab27A* CRISPR-knockout cells: clones 1, 2 and 3 (KO#1, KO#2 and KO#3). Data was generated from 3 independent experiments. **(E)** Percentages of cells within each stage of the cell cycle of WM164 cells expressing the non-targeting shRNA control, *RAB27A* shRNA#1 or *RAB27A* shRNA #2. Error bars represent the mean ± SEM of 3 independent experiments. Non-targeting control was compared to the other samples via ANOVA and a Dunnett’s test and no significance was detected. **(F)** Left, cell counts over the indicated hours in ectopically RAB27A-overexpressing C8161 cells. Right, Western blotting showing the levels of endogenous RAB27A (∼25kDa) and overexpressed RAB27A (RAB27A-GFP, ∼50kDa). NS, not significant.

To further validate that the effect of abrogating RAB27A expression on proliferation is only transient, we carried out a complete knockout of *Rab27A* in B16-F10 melanoma cells employing CRISPR-Cas9 technology (Figure 3D). B16-F10 is a very well-characterized murine melanoma cell line with a high expression of RAB27A (Figure 1A, B). Single cell clones of *Rab27A* knockout B16-F10 were generated and assessed for proliferation. Similar to WM164 and WM983C, no significant differences in proliferation post *Rab27A* KO were observed (Figure 3D). However, the capacity of B16-F10 cells to metastasize had significantly reduced post *Rab27A* KO *in vivo* as reported in our previous study (Guo, Lui, et al., 2019).

Recent reports have linked RAB27A with the regulation of cell cycle genes in other cancer types (Feng et al., 2017; Liu et al., 2017). Therefore, we carried out *RAB27A*-KD in FUCCI (Fluorescent Ubiquitination-based Cell Cycle Indicator) expressing WM164 cells (Haass et al., 2014; Spoerri, Beaumont, Anfosso, & Haass, 2017) and examined the proportion of cells within each phase of the cell cycle. Interestingly, we found no statistically significant difference in the cell cycle profile between control and *RAB27A*-knockdown cells (Figure 3E), which suggested that the transient proliferation defect caused by *RAB27A*-knockdown was not related to skewing of the cell cycle.

Finally, we carried out overexpression of RAB27A in RAB27A low cell line C8161. Overexpression of RAB27A did not cause any change in the cellular proliferation of C8161 cells suggesting that C8161 proliferation is indeed independent of RAB27A function (Figure 3F).

Therefore, in this study, we demonstrate that *RAB27A*-knockdown impacts cell proliferation and colony formation potential and has a modest effect on survival in metastatic melanoma cell lines that express high levels of RAB27A. However, in contrast to the persistent reduction in cell invasion and metastasis post abrogating *RAB27A* expression as reported previously (Guo, Lui, et al., 2019), we find that all melanoma cell lines tested regained proliferative potential over time. Here, we offer an explanation for the apparent contradiction to previous studies by others demonstrating that abrogating *RAB27A* expression in melanoma and other cancer cells reduces cell proliferation and colony formation (Akavia et al., 2010; Bobrie et al., 2012; Li et al., 2019; Li et al., 2017; Liu et al., 2017). The possible reason for these differences could be due to variations in the employed methodologies. Two studies used siRNA-based transient transfection to demonstrate that knocking down RAB27A induced modest increase of apoptosis, enhanced sensitivity to cisplatin and reduced the invasive and proliferative capacity of pancreatic (Li et al., 2017) or bladder cancer cells (Liu et al., 2017). An elegant study demonstrated that shRNA-expressing lentiviral mediated RAB27A blockade in breast cancer cells (mammary carcinoma 4T1) led to a reduced secretion of exosomes and reduced the primary tumor growth as well as metastasis to lungs. However, the authors clearly state that they used these cells within a one-month period post-lentiviral transduction (Bobrie et al., 2012). The seminal study, which identified RAB27A as a tumor dependency gene in melanoma, also demonstrated reduced proliferation post shRNA mediated knockdown of RAB27A in RAB27A-high cells (Akavia et al., 2010). However, they also looked at earlier time-points as they measured growth till day 6 post-knockdown. In addition, they found changes in cell cycle genes after *RAB27A*-knockdown (Akavia et al., 2010). Notably, we also observed *CDK2* as a gene associated with RAB27A-high melanoma clinical samples in our recent study (Guo, Lui, et al., 2019). However, here we discovered that the cell cycle profile did not change significantly after *RAB27A*-knockdown. In summary, most of these studies use *RAB27A*-abrogated cells before the compensation that we describe here has fully taken effect. It is reasonable to use freshly transduced/transfected cells to ensure efficient knockdown of *RAB27A*, but in our study, we confirmed the persistent *RAB27A* knockdown after culturing the cells for multiple passages (more than a month) and discovered the loss of proliferation defect. Moreover, we confirmed these findings by CRISPR-mediated knockout of *Rab27a* in murine melanoma cells. This proves the existence of a compensation mechanism specifically for the proliferation defect rather than simply loss of RAB27A knockdown over time. We hypothesized that the proliferation defect post RAB27A downregulation is mediated through the upregulation of another RAB-family member. However, in our study, the closely related RAB27B did not appear to be responsible for the rescue of the proliferation defect. RAB27A is transcriptionally activated by Microphthalmia-associated Transcription Factor (MITF) (Garraway et al., 2005). In another study, we have analyzed proteomics data of melanoma cells expressing endogenously high versus low levels of MITF and RAB27A; we have also compared MITF-knockdown and ectopically MITF-overexpressing cells to their wild-type controls (Haass laboratory, unpublished data). Interestingly, this study revealed that in MITF^low^ cells, RAB27A is downregulated but RAB11A, RAB7A and RAB2A appear upregulated. Accordingly, in MITF^high^ cells, the reverse is true. This finding indicates that not RAB27B but RAB11A, RAB7A and/or RAB2A could potentially compensate for the loss of RAB27A. However, this observation will require further assessment in future studies. Importantly, the defects in cell invasion and metastasis post-abrogating RAB27A expression, also seen in the studies discussed above, were found to be persistent (Guo, Lui, et al., 2019).

In conclusion, although RAB27A is an important promoter of melanoma invasion and metastasis, our study demonstrates that its effects on cell proliferation and survival, in contrast to migration, invasion and metastasis (Guo, Lui, et al., 2019), are not persistent. This finding provides a deeper understanding of RAB27A function in melanoma cell biology.

## Acknowledgements

This work was supported by project grants to K.A.B. (1051996, Priority-driven collaborative cancer research scheme/Cancer Australia/Cure Cancer Australia Foundation), to N.K.H. (RG 13-06, Cancer Council New South Wales; APP1003637 and APP1084893, National Health and Medical Research Council) and to ST (Tour De Cure, Pioneering Research Grant).

## Supplementary Material to

### Materials and Methods

#### Cell lines

The human melanoma cell lines C8161, 1205Lu, WM164 and WM983C were genotypically characterized (Davies et al., 2009, Smalley et al., 2007a, Smalley et al., 2007b) and grown as described (Spoerri et al., 2017). Cell line authentication was accomplished by STR fingerprinting (Molecular Genetics facility, Garvan Institute of Medical Research, Darlinghurst, NSW, Australia). Stable expression of Fluorescence Ubiquitination Cell Cycle Indicator (FUCCI) in WM164 cells was performed as previously described (Haass et al., 2014, Spoerri et al., 2017). Mycoplasma contamination in all cell lines was ruled out via PCR profiling at the Garvan Institute of Medical Research.

**Lentiviral transductions** were performed as previously described (Guo et al., 2019). Briefly, *RAB27A* shRNA or non-target shRNA lentivirus particles (MISSION, Sigma/Merck) were produced using the calcium phosphate precipitation method in HEK293T cells. Melanoma cell lines were transduced with lentivirus and 4µg/mL polybrene followed by selection with puromycin (2µg/mL).

#### Proliferation assays

Cells were seeded at a density of 20,000 cells/well in 12-well plates (triplicate wells per condition), trypsinized and counted by hemocytometer on days 1-4 after seeding.

#### Rab27A CRISPR

Rab27A CRISPR knockout was performed as previously described (Guo et al., 2019). Briefly, guide RNAs targeting *Rab27A* were designed as described (Ran et al., 2013) and cloned into the pX330-IRES-GFP vector. Melanoma cell lines were transfected with the vector alone and plasmids encoding *Rab27A* gRNA. Post-transfection, GFP positive cells were FACS sorted. Single cell cloning was then achieved by serial dilution. The clones were expanded and *Rab27A* knockout was confirmed by Western blotting.

#### Colony-formation assays

2000 cells (duplicate wells per condition) were seeded in 12-well plates, with culture medium changed twice weekly. After 2 weeks of growth, colonies were gently washed with PBS, fixed on ice with methanol for 15 min, and then stained with 0.5% crystal violet/methanol for 15 min. Excess stain was removed by carefully washing with distilled water. Plates were left to dry at RT for 1h and then scanned using a color scanner (Konica Minolta Bizhub C360). Quantification of colony formation was determined by percentages of stained area/total area per well using ImageJ software.

**Western blotting** was performed as previously described (Guo et al., 2019). The following antibodies were used: mouse anti-human RAB27A (WH0005873M2, Sigma-Aldrich, Merck KGaA), rat anti-tubulin (Ab6160, Abcam, Cambridge, MA, USA). Membranes were washed with TBS-T for 30min, then probed with the appropriate secondary HRP conjugated antibody (Sigma) for 1-2h at room temperature. Proteins were visualized with SuperSignal West Pico chemiluminescent substrate (ThermoFisher Scientific) and the ChemiDoc MP System (BioRad).

## Figure Legends

**Supplementary Figure 1.**
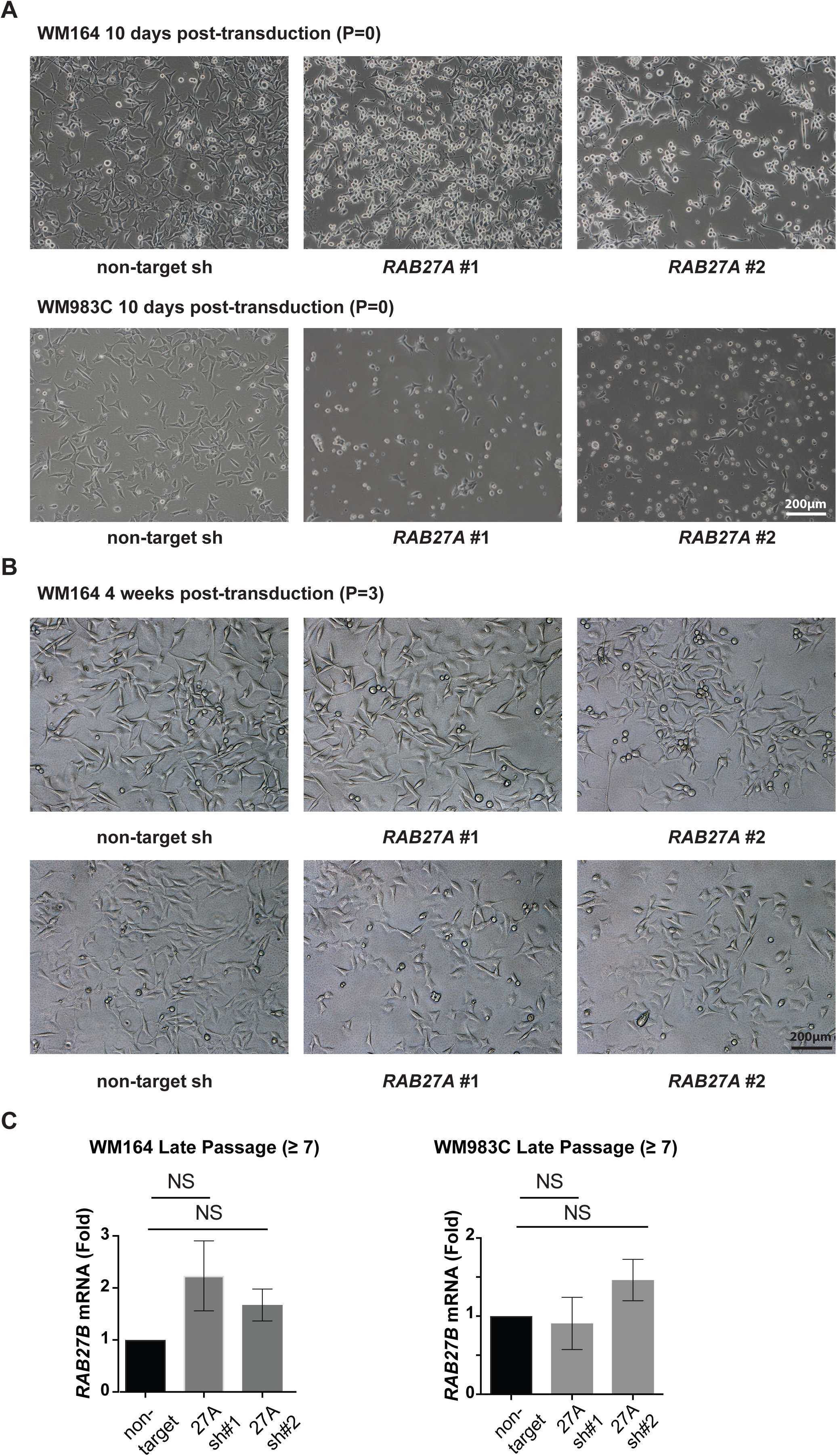
**(A)** High magnification phase contrast images of WM164 and WM983C melanoma cells approximately 10 days after transduction with *RAB27A* shRNA or control lentivirus and 8 days after selection with puromycin. *RAB27A* shRNA KD cells are at Passage 0 after transduction. Scale bars = 200μm. Representative of 3 independent experiments. **(B)** High magnification phase contrast images of WM164 and WM983C melanoma cells approximately 4 weeks (3 passages) after transduction with *RAB27A* shRNA or control lentivirus Scale bars = 200μm. Representative of 3 independent experiments. **(C)** Real time PCR TaqMan analysis of *RAB27B* mRNA expression in WM164 and WM983C cells (≥ 7 passage) transduced with the indicated *RAB27A* shRNA or control constructs. Error bars represent the mean ± SEM of 3 independent experiments. Non-targeting control was compared to the other samples via ANOVA and a Dunnett’s test and no significance was detected. NS, not significant.

